# predicTTE: An accessible and optimal tool for time-to-event prediction in neurological diseases

**DOI:** 10.1101/2024.07.20.604416

**Authors:** Marcel Weinreich, Harry McDonough, Nancy Yacovzada, Iddo Magen, Yahel Cohen, Calum Harvey, Sarah Gornall, Sarah Boddy, James Alix, Nima Mohseni, Julian M Kurz, Kevin P Kenna, Sai Zhang, Alfredo Iacoangeli, Ahmad Al-Khleifat, Michael P Snyder, Esther Hobson, Ammar Al-Chalabi, Eran Hornstein, Eran Elhaik, Pamela J Shaw, Christopher McDermott, Johnathan Cooper-Knock

## Abstract

Time-to-event prediction is a key task for biological discovery, experimental medicine, and clinical care. This is particularly true for neurological diseases where development of reliable biomarkers is often limited by difficulty visualising and sampling relevant cell and molecular pathobiology. To date, much work has relied on Cox regression because of ease-of-use, despite evidence that this model includes incorrect assumptions. We have implemented a set of deep learning and spline models for time-to-event modelling within a fully customizable ‘app’ and accompanying online portal, both of which can be used for any time-to-event analysis in any disease by a non-expert user. Our online portal includes capacity for end-users including patients, Neurology clinicians, and researchers, to access and perform predictions using a trained model, and to contribute new data for model improvement, all within a data-secure environment. We demonstrate a pipeline for use of our app with three use-cases including imputation of missing data, hyperparameter tuning, model training and independent validation. We show that predictions are optimal for use in downstream applications such as genetic discovery, biomarker interpretation, and personalised choice of medication. We demonstrate the efficiency of an ensemble configuration, including focused training of a deep learning model. We have optimised a pipeline for imputation of missing data in combination with time-to-event prediction models. Overall, we provide a powerful and accessible tool to develop, access and share time-to-event prediction models; all software and tutorials are available at www.predictte.org.

## Research in Context

### Evidence before this study

Predicting time-to-event (e.g. survival) is an important goal across almost all human diseases. We reviewed journal articles describing methods for ‘time-to-event’ and, particularly ‘survival’ prediction. We focused on applications in ‘amyotrophic lateral sclerosis’ (ALS) where we are well-placed to assess the extent to which prediction models are used in the clinic. Regarding imputation of missing data we searched journal articles pertaining to the use of ‘machine learning’; we identified the ‘MissForrest’ random-forest model together with several use-cases demonstrating efficacy. For time-to-event prediction, deep learning models within the ‘pycox’ package, which is a python implementation of PyTorch based time-to-event models, were identified as likely to have the best performance. However, the application of these models in clinical scenarios was limited or absent. We also identified the ‘flexsurv’ package, which is an R implementation of a range of parametric models for time-to-event tasks, as a complementary set of models which could provide comparable or even superior performance to deep learning models, particularly when the training dataset is smaller.

### Added value of this study

Time-to-event prediction of disease milestones such as onset and survival are essential to guide clinical practice and personalised treatment choices. Yet, research is often hindered by a gap between clinicians with access to data, and researchers who possess optimal tools. Our mixed team used clinical data to develop predicTTE, an integrated pipeline which enables a researcher with a dataset but no bioinformatics experience, to design an optimal prediction model and make it available via an online portal to end-users. In addition we provide the capacity for end-users to directly contribute additional training data. We anticipate a positive-feedback loop whereby the performance of trained models is improved progressively via increases in training data, which is sourced directly from the community of clinicians and patients, while maintaining patient privacy and offering appropriate credit.

predicTTE consists of a customizable ‘app’ and accompanying online portal for any time-to-event analysis. It implements state-of-the-art models, including all the deep learning models and parametric spline approaches from the ‘pycox’ and ‘flexsurv’ packages, respectively. We provide the capacity to develop ensemble models where a second round of focused training is performed using a subset of the most informative training data, based on an initial prediction. We also implement ‘MissForrest’ for missing data imputation, and demonstrate the optimum form of this model using an independent validation cohort. Our work will inform future practice in this field.

### Implications of all the available evidence

We provided use-cases demonstrating how our pipeline can be used for survival prediction, biomarker assessment and personalised medicine in three different neurological disease scenarios. We combine optimum time-to-event prediction tools in an accessible form which facilitates efficient data sharing. Going forward, our work could be adapted to include additional models and datasets.

## Introduction

Understanding determinants of time-to-event is an important problem with relevance in any clinical context including longitudinal progression. Neurological disease has suffered a dearth of predictive biomarkers, in part because of difficulty accessing the site of disease within the nervous system^1^. An alternative to better markers of molecular pathogenesis is to make better use of available data within a suitable model. Cox regression^2^ is a popular model for time-to-event tasks which assumes a fixed proportional-hazard ratio, whereby the relative hazard-rate between patients is invariable over time. This is an unrealistic assumption for many contexts and has likely led to misinterpretation of the underlying drivers of time-to-event. Many new models have been developed (e.g. using deep learning^3^) but usability is limited by requirement for computational expertise. In the clinical context this often excludes important users such as clinicians and patients. We address this limitation in a new app and accompanying online platform where we implement a range of cutting-edge models for model design, hyperparameter tuning and prediction. The online platform includes the capacity to provide end-users such as patients and clinicians with secure access to a trained model for prediction, and with the option to contribute new data. We anticipate that this facility could rapidly scale to provide largest-in-class datasets and optimal prediction performance. We have named our software ‘predicTTE’ (predicting time-to-event); our implementation is summarised in **Fig. 1**. Here we demonstrate three use-cases from example Neurological diseases: two focused on prediction of survival in amyotrophic lateral sclerosis (ALS) using clinical and/or biofluid biomarkers; and thirdly prediction of time to all-cause mortality for patients treated with anticoagulation for atrial fibrillation (AF). We show how predicTTE can be used to provide clinical predictions, to evaluate the relative value of biomarkers, and to assign an optimal drug treatment in a personalised manner. ALS is an exemplar because of the lack of a sensitive and specific biomarker, and the fact that the key cellular pathology within motor neurons (MN), is accessible only after death.

**Figure 1:**
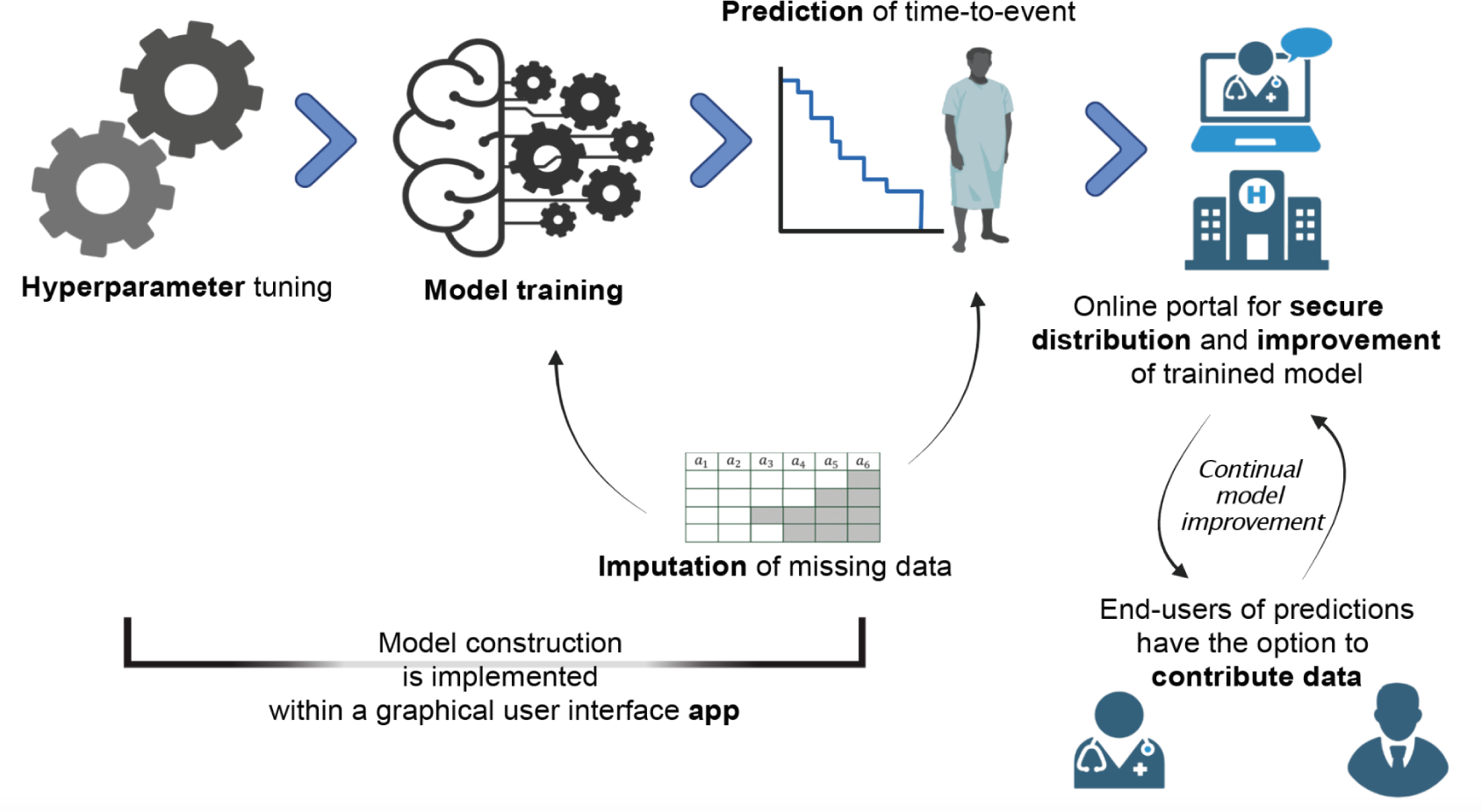
Accessible and optimal time-to-event prediction is achieved through implementation of cutting-edge models within an app and online platform. We have implemented a set of deep learning and spline models for time-to-event modelling, including model selection, hyperparameter tuning, model training and imputation of missing data. This functionality is provided within a fully customizable ‘app’ (left panels). An accompanying online portal includes capacity for end-users including patients, clinicians and researchers, to access and perform predictions using a trained model, and to contribute new data for model improvement, all within a data-secure environment (right panels). All software and tutorials are available at www.predictte.org.

A key aspect of our platform is the capability to impute missing data using a random forrest model called MissForest which has shown superior performance in real-world testing.^4,5^ Missing data is a common real-world problem and failure to handle missing data can limit usability. We develop and test an implementation of MissForrest for combination with deep learning prediction models; we show that our implementation can handle missing data in >50% of training instances without significantly degrading prediction performance, and is optimal for use in downstream applications such as discovery of genetic drivers of disease survival.

## Results

The capabilities and potential of predicTTE are demonstrated in three Neurological disease use-cases.

### Use-case 1: Predicting survival in ALS, optimisation of model training and imputation of missing data

Amyotrophic lateral sclerosis (ALS) is an archetypal neurodegenerative disease associated with irreversible progression. For most patients, it leads to death within 2 to 5 years, primarily due to respiratory failure.^6^ Importantly ALS survival is thought to be largely a function of disease progression i.e. death does not usually occur from an unrelated cause. Given that the disease progression is observable at diagnosis, we and others^7^ have hypothesised that accurate survival prediction is feasible based on baseline clinical measurements.

We chose a set of baseline clinical variables with evidence for a relationship with ALS survival, and with previous evidence in survival prediction:^7^ age, presence/absence of an ALS-associated *C9ORF72* mutation,^8^ site of disease onset, diagnostic delay, ALSFRS-R slope, El-Escorial category,^9^ and presence/absence of frontotemporal dementia (FTD). Diagnostic delay is the time from symptom onset to diagnosis with ALS and has been consistently linked to ALS survival,^10^ probably because it represents the speed of progression to the point where the disease is both clinically manifest and the patient has sufficient functional impairment to seek medical assistance. The ALSFRS-R is a commonly used functional rating scale for ALS^11^; to infer the rate of change or ‘slope’ we assumed a linear decline between the time of symptom onset and the time of diagnosis i.e. over the period which constitutes the diagnostic delay. Importantly all of these data are frequently collected at ALS diagnosis, including the El-Escorial category, which is calculated from routine neurophysiological assessment. We also added sex because there is consistent evidence that sex impacts ALS biology^12^ and the data are typically available.

Hyperparameter tuning (**Supplementary Fig. 1, Methods**) selected a PCHazard deep learning model^13^ with parameters detailed in **Supplementary Table 1**. Training utilised 5,336 ALS patients from Project MinE^14^ who were recruited from ALS clinics throughout Europe (**Methods**); for 4,053 of these patients an accurate survival time was recorded because the patient was deceased, and for the remainder we relied upon a censored survival time. We trialled an ensemble approach based on the hypothesis that patients with similar predicted survival are more informative regarding the actual survival of a test patient and similarly, patients with very different survival may be relatively uninformative (**Fig. 2a**). Our ensemble model works as follows: In the first step we applied a trained model to generate a prediction for the exact time point when a patient is more likely to have died than to be surviving. In the second step the model is further trained before delivering a final prediction; but this training uses only a subset of the training cohort including individuals with measured survival within a range defined by the initial prediction (**Methods, Fig. 2b-c**).

**Figure 2:**
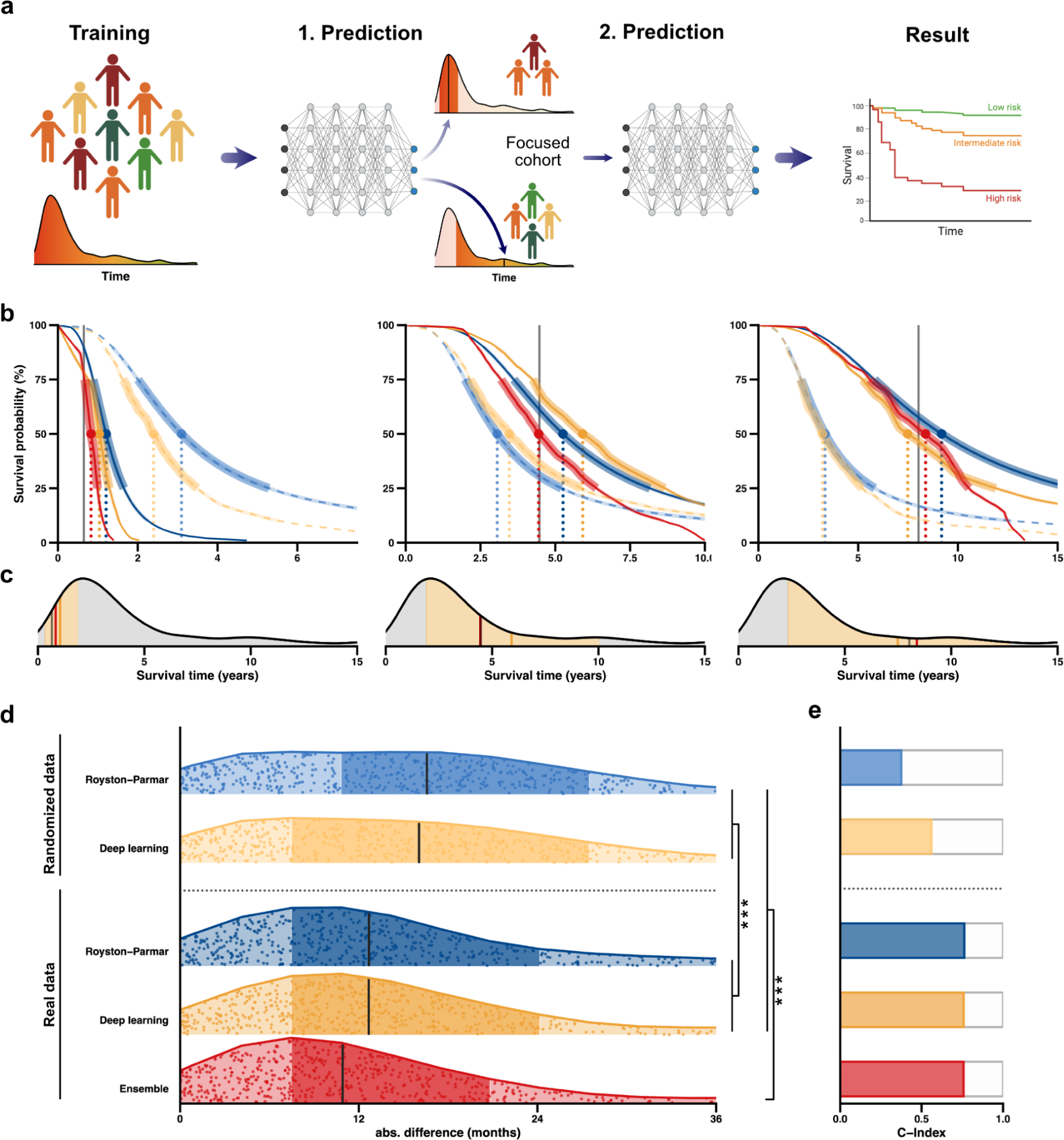
Use-case 1: Use of predicTTE to predict survival in ALS. (**a)** Ensemble model structure consisting of two stages of PCHazard model training and prediction. Initially the prediction model is trained using the entire training cohort and a prediction of median survival is outputted for the test patient; in the second stage further training is applied using a focused cohort of patients most similar to the predicted survival from the first stage (**Methods**). The final prediction of survival time for the test patient is the output from the model following the second stage of training. (**b-c**) Illustrations of survival probability outputs for the various models, for example patients with survival <2 years, 2-5 years and 5-10 years. Dotted lines indicate randomised training data for comparison with the real training data (solid lines). Blue indicates the optimum spine model, yellow indicates the optimum deep learning model and red indicates the ensemble model. In (**c**) the total set of patients utilised in stage one of ensemble training is indicated by grey shading and the observed time by the grey line; the focused cohort used in the second stage of ensemble model training is indicated by yellow shading with the yellow line being the initial prediction and the red line indicating the final prediction. (**d-e**) Accuracy of survival prediction in the UK validation cohort using models trained in the Project MinE cohort. (**d**) Absolute difference between predicted and observed survival is plotted for each individual; vertical lines indicate median values, darker shading indicate the 25-75% quantile range. P-values are shown for a Wilcoxon rank-sum test comparing absolute difference between predicted and observed survival for each model. ***p<0.01. (**e**) C-index for model performance; models are as specified in (**d**).

Testing was performed on an entirely independent UK validation cohort (n=661 ALS patients including 595 patients with observed survival, **Methods**). The ensemble model delivered a median absolute difference between predicted and actual survival of 10.9 months (se=1.4 months) (**Fig. 2b, Supplementary Table 2**) and C-index (concordance, **Methods**) of 0.76 (**Fig. 2e, Supplementary Table 2**). The ensemble approach significantly improved prediction accuracy compared to the optimum spline model – Royston Palmar – and to a single stage of training of the PCHazard model (t-test, p<0.01, **Fig. 2d, Supplementary Table 2, Supplementary Fig. 3**). Concordance was not significantly improved by model choice (**Fig. 2e, Supplementary Table 2**). As an additional comparison we repeated hyperparameter tuning and model training using training data with randomly shuffled clinical variables (**Methods**); as expected prediction performance in the UK validation cohort was significantly impaired (**Fig. 2b-e)**.

### Data missingness is frequent and can be imputed using MissForest

Reported results include imputation of missing covariates via a random-forest model called MissForest.^4^ Missing data is a common real-world phenomenon, which necessitates the adoption of imputation methods that can yield highly accurate results. Moreover, relevant for this particular use-case, the survival profile of ALS patients with missing data is not equivalent to those without missing data^15^ and thereby, a model that neglects patients with missing data does not capture the full range of ALS phenotypic variation. Numbers and proportions of missing data used in training are detailed in **Supplementary Table 3**.

MissForest was trained using the relationships between observed covariates in the training dataset. To test the performance of the MissForest model we randomly selected and omitted 50 rounds of 50 data points from each covariate in the training dataset (**Methods**). These data were then imputed using the MissForest model and the correlation between imputed and correct values was calculated. For continuous variables we demonstrate that MissForest achieves a statistically significant correlation between actual and imputed values (age of onset: Pearson correlation, r=0.27, p=8.61e-17; diagnostic delay: *r*=0.31, *p*=9.35e-49; ALSFRS-R slope: *r*=0.53, *p*=3.12e-137; **Fig. 3a, c**), supporting the efficacy of this imputation strategy.

**Figure 3:**
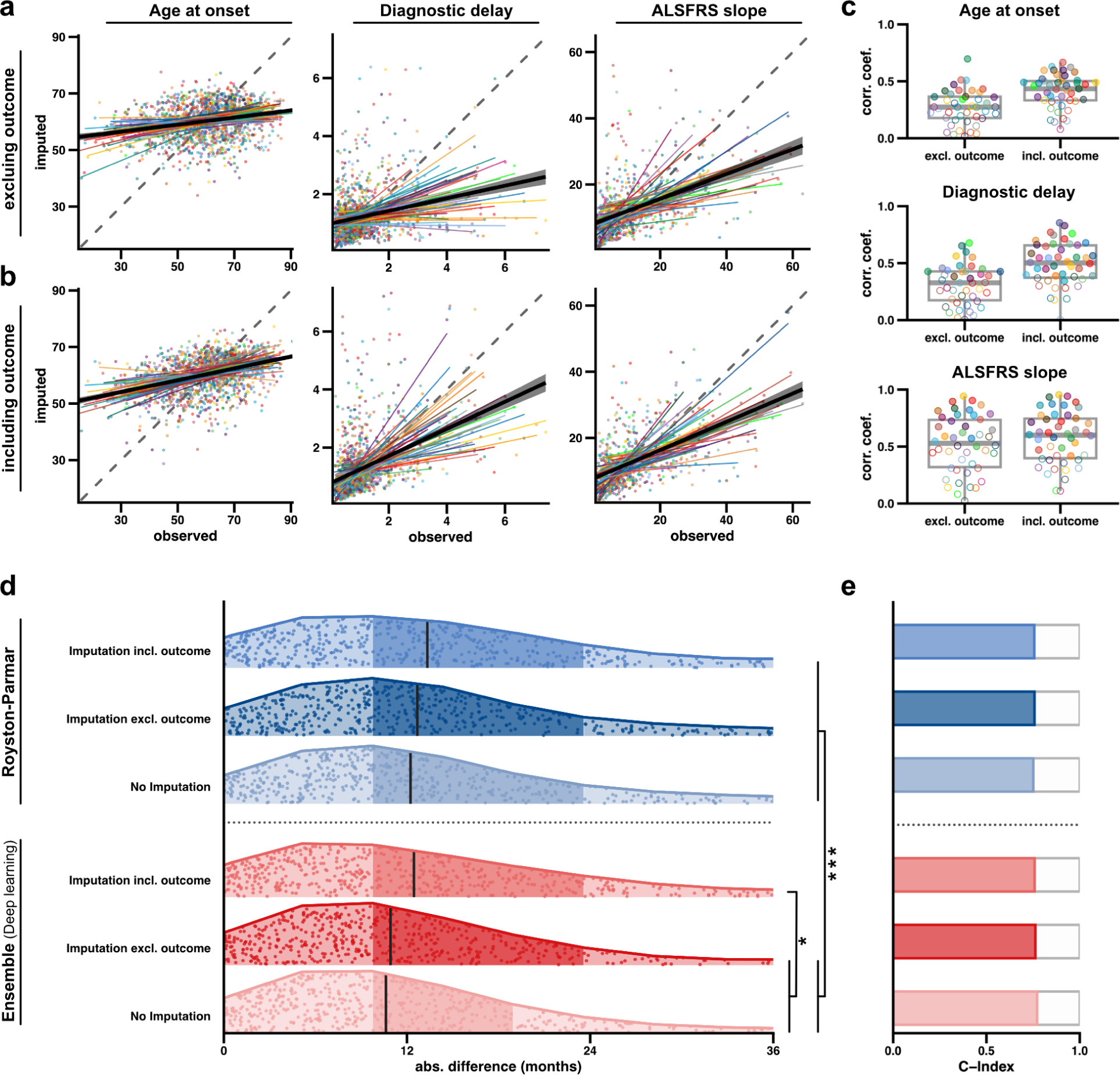
Missing datapoints can be imputed using MissForest without degrading prediction performance. To evaluate MissForest^4^ the model was trained and used to impute held-out covariate values within the training dataset. We randomly selected and omitted 50 rounds of 50 data points from each covariate in the Project MinE training dataset. Results are shown for ALSFRS-R-slope, diagnostic delay, and age of onset. Imputation was performed (**a**) not including and (**b**) including the outcome variable (ALS survival). (**a-b**) Each point in the plot represents a single instance of imputation for an observed covariate. The coloured lines depict the median linear regression fits to the imputed values for each round, and every colour indicates a different round. The black line represents the median linear regression fit for all imputed values from all 50 rounds of imputation. This visualisation effectively demonstrates the stability and consistency of the prediction performance. The correlation coefficients are summarised in (**c**), where the filled dots represent significant correlations (Pearson correlation, p<0.05). As expected, overall the imputation performance is better if the outcome is included. (**d-e**) Accuracy of survival prediction using models trained in the Project MinE cohort; assessment is performed in an independent UK validation cohort including 661 ALS patients where survival time was observed in n=595. Accuracy is shown for survival prediction models with and without imputation of missing data; and where imputation is performed with and without inclusion of the outcome variable. (**d**) Absolute difference between predicted and observed survival is plotted for each individual; vertical lines indicate median values, darker shading indicate the 25-75% quantile range. P-values are shown for a Wilcoxon rank-sum test comparing absolute difference between predicted and observed survival for each model. * p<0.05; ***p<0.01. (**e**) C-index for model performance; models are as specified in (**d**).

An important decision in imputation of missing data is whether to include the outcome variable in imputation. Inclusion can increase the risk of overfitting to the training set whereas omission can artificially depress the importance of imputed data points.^16^ However, if the missingness of the covariate is a determinant of the relationship between the covariate and the outcome variable (e.g. if short survivors are less likely to perform all tests) then imputing including the outcome variable relies on an incorrect assumption.^17^ To evaluate these alternatives we performed imputation, training and prediction under both scenarios: with and without the use of survival in the imputation of missing datapoints; and also using a training dataset without any imputed data (n=1,683) (**Methods**). As expected, imputation performance improved with inclusion of the outcome variable (**Fig. 3b-c**). However, despite this, prediction performance in the UK validation cohort was most accurate using the ensemble model where imputation of missing data was performed *without* the outcome variable (**Fig. 3d, Supplementary Table 2, Supplementary Fig. 3**). Inclusion of the outcome variable in imputation reduced the accuracy of prediction in the validation cohort for both the ensemble model and the optimum spline model (**Fig. 3d, Supplementary Table 2, Supplementary Fig. 3**). As expected, prediction accuracy for patients with complete data, i.e. where imputation was not necessary, was better than for more inclusive models trained with imputed data; however this difference was relatively small and not statistically significant (**Fig. 3d, Supplementary Table 2, Supplementary Fig. 3**). We conclude that the prediction model we have implemented can accommodate realistic clinical data including missing covariates, without significant decline in performance.

### Predicted survival times can be used for genetic discovery

In addition to its clinical importance, ALS survival is also an important endpoint for discovery of the molecular drivers of pathogenesis. We hypothesised that predicted survival could be used in genetic discovery when actual survival was not available. rs12608932 within *UNC13A* is a validated genetic modifier of ALS survival.^18^ For 5,498 patients from Project MinE (www.projectmine.com) we first predicted survival using our optimum ensemble model, and then used Cox regression with platform and first 10 PCs as covariates to test the effect of the rs12608932(C) allele on predicted survival. This analysis used only predicted and not measured survival; indeed for many of these patients actual survival data was unavailable. As expected^18^ there was a significant negative impact of the rs12608932(C) allele on predicted survival (coef=+0.05, p=0.02, Cox Regression). This suggests that predicTTE predictions could be used to improve discovery of molecular mechanisms underlying ALS survival particularly in cohorts where longitudinal phenotypic information is not available. By the same logic, predictions could be used to stratify clinical trial patients based on a baseline assessment.

Notably, using survival predictions where outcome *was* included in the imputation of missing data, did not result in a significant link between UNC13A genotype and ALS survival (coef=+0.04, p=0.09). This supports our conclusion that the outcome variable should not be used in imputation of missing data in conjunction with our prediction model.

### Use-case 2: Survival prediction to evaluate candidate biomarkers in ALS

In use-case 1, our focus was on predicting the time to an outcome variable: ALS survival. Another major use of an optimum time-to-event prediction model is in the evaluation of specific covariates, such as a candidate biomarker, and their predictive performance. This is a strength of the Cox model^2^ where hazard ratios are recorded for each covariate. In contrast, an equivalent covariate-specific measure is not reported in our pipeline. However, in this use-case we show how specific covariates can be evaluated using predicTTE in a manner which exploits the enhanced accuracy gained from using an optimal model.

In our previous case study of ALS, we focused on the use of baseline clinical data to predict ALS survival. The use of blood-based biomarkers has recently revolutionised the field^19,20^ and was key to the recent FDA approval of Tofersen to treat ALS caused by mutations within *SOD1.*^21^ We applied predicTTE to a previously published dataset^19^ to evaluate how survival prediction using clinical symptoms is improved by inclusion of two biomarkers: plasma concentration of NfL (neurofilament light chain) and/or plasma levels of mir181 (**Fig. 4a**).

**Figure 4:**
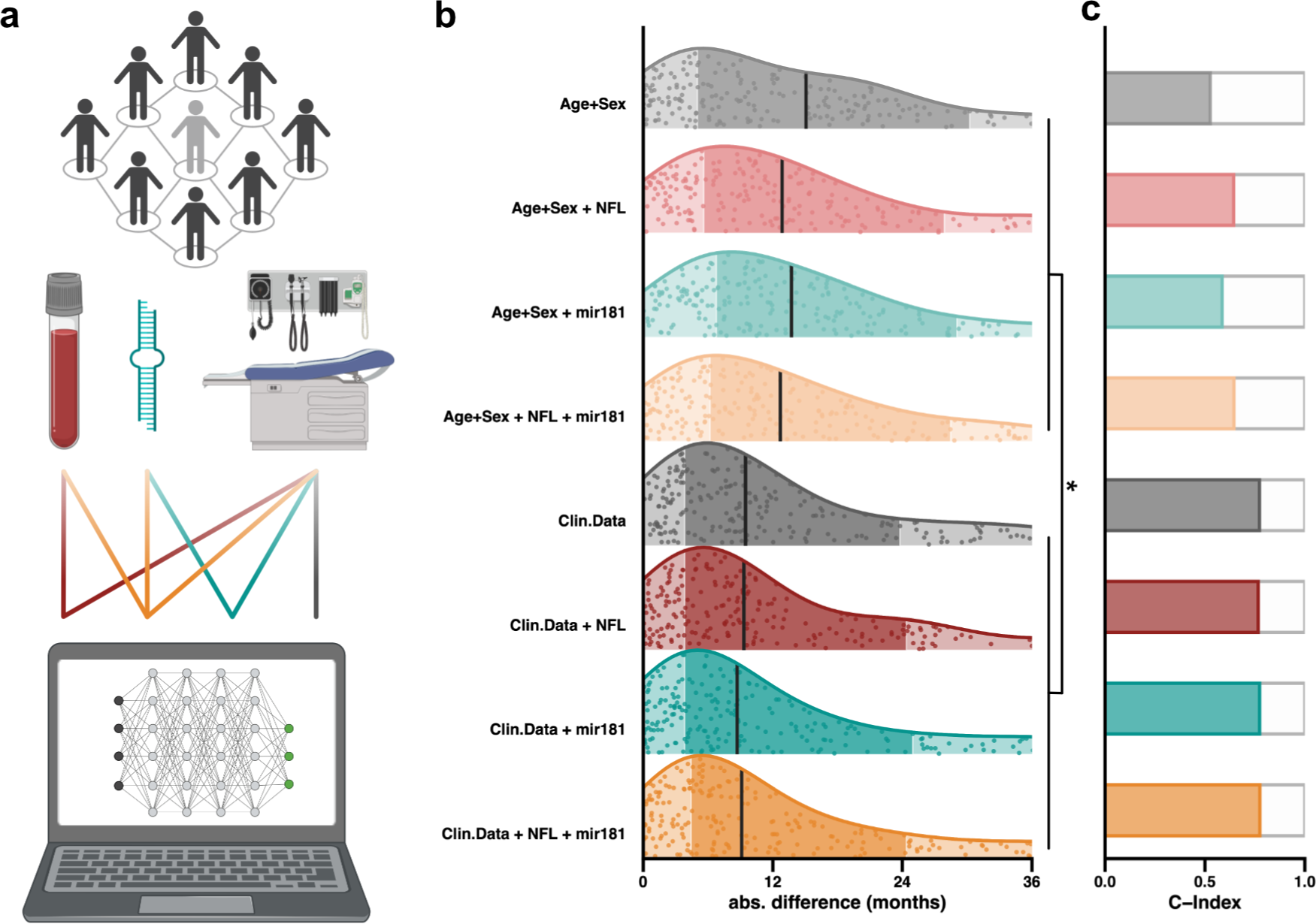
Use-case 2: Survival prediction to evaluate candidate biomarkers in ALS. (**a**) After hyperparameter tuning and training, the optimal model (here PCHazard) was trained and used to predict ALS survival. The set of model covariates, including clinical features and blood-based biomarkers, was varied to allow evaluation of each covariate as a determinant of model performance. (b) Absolute difference between predicted and observed survival is plotted for each individual for each of trained model; vertical lines indicate median values, darker shading indicate the 25-75% quantile range. P-values are shown for a Wilcoxon rank-sum test comparing absolute difference between predicted and observed survival for each model. * p<0.05. As measured by either model median absolute difference between observed and predicted survival (**b**) or concordance (**c**) the best prediction performance is achieved including clinical features plus plasma concentration of mir181.

Unlike the first use-case, here clinical measurements were not performed at baseline. Therefore, we have included ALSFRS-R at the time of observation in addition to the ALSFRS-R slope. Other clinical covariates included were: age, sex, site of disease onset, and time from symptom onset to time of evaluation. We have predicted survival from onset of symptoms, exactly as in the first use-case. For n=248 patients, missing data was restricted to only 5 plasma concentrations of NfL. This use-case did not include an independent validation cohort and therefore we performed hyperparameter tuning and evaluated model performance in the same cohort via nested cross-validation (**Methods**). During hyperparameter tuning again the PCHazard model^13^ performed best (**Supplementary Table 1**), an ensemble model was not evaluated because of the small sample size.

Comparison of prediction performance for models including all covariates or clinical characteristics with and without plasma biomarkers revealed that, whether performance is measured by C-index (**Fig. 4c, Supplementary Table 4**) or by median absolute difference in predicted versus observed survival (**Fig. 4b, Supplementary Table 4**), the best performance was achieved when plasma mir181 was combined with clinical measures for survival prediction. The largest improvement in model performance was achieved by incorporating clinical variables including ALSFRS-R (slope and value at time of assessment) and site of disease onset; but prediction performance using blood based biomarkers plus age and sex was significantly better than for age and sex in isolation (Wilcoxon rank-sum test, p<0.05, **Methods, Fig. 4b**). We conclude that both plasma NfL and particularly mir181 contribute independent information to the prediction performance (**Fig. 4b**) and should be evaluated in a larger cohort.

We have demonstrated that predicTTE could also be used to evaluate prediction performance for selected covariates in subsets of data and even in specific individuals; this is a key step towards personalised medicine which is the focus of use-case 3.

### Use-case 3: Individualised prediction of all-cause mortality to guide preventative drug treatment

Cerebrovascular stroke is a potentially fatal complication of atrial fibrillation (AF) due to cardiac thromboembolism. Current practice is to initiate anticoagulation with a direct oral anticoagulant (DOAC), except in cases where there is excessive risk of haemorrhage. However, with several options available, the choice of DOAC is guided by grouping of patients based on age and comorbidities such as renal impairment.^22^ This practice is likely to be suboptimal for certain individuals; an ideal system would predict outcome *for each individual* based on their personal characteristics. Currently randomised clinical trials have focused on relatively homogenous cohorts and fail to provide evidence for many diverse individuals.^23^ Here we apply predicTTE to a previously published dataset describing clinical characteristics, DOAC treatment and outcome measures in patients with AF (**Fig. 5a**).^24^

**Figure 5:**
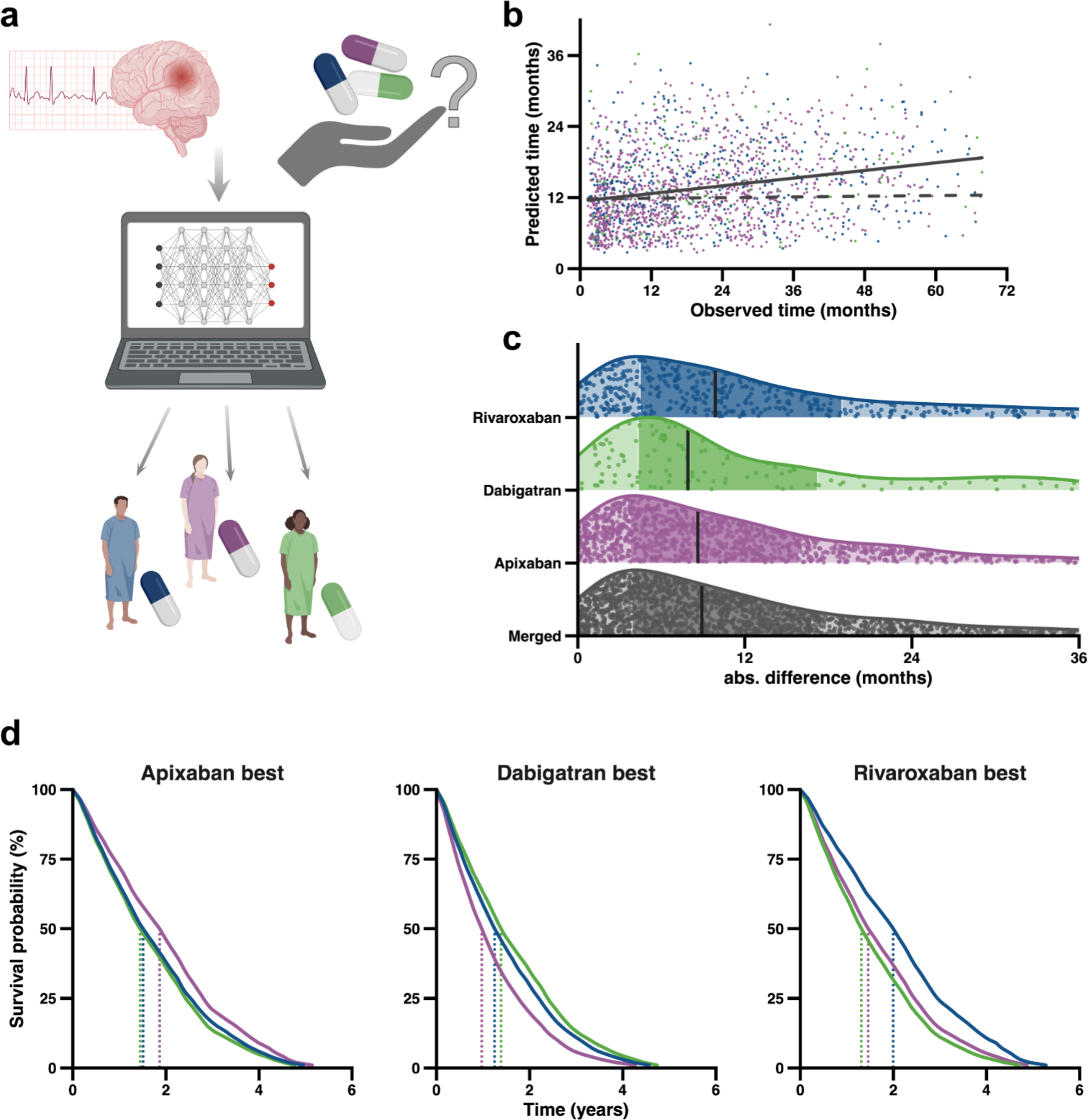
Use-case 3: Individualised prediction of all-cause mortality to guide preventative drug treatment. (**a**) After hyperparameter tuning and training, the optimal (here PCHazard) model was used to predict time to all-cause mortality after diagnosis of AF post-stroke. (**b**) Comparison of observed and predicted time to all-cause mortality (coloured by DOAC) together with line of best fit demonstrating a positive correlation (solid line), compared to predictions made using a model trained with randomised covariates (dashed line). (**c**) Absolute difference between predicted and observed survival is plotted for each individual; vertical lines indicate median values, darker shading indicate the 25-75% quantile range. (**d**) Three example patients with varying clinical characteristics, demonstrating how choice of anticoagulation with either apixaban (purple), dabigatran (green), or rivaroxaban (blue), impacts predicted survival time. For each DOAC the patient that has the best predicted outcome is shown in comparison to the predicted outcome using the other 2 DOACs. Clinical characteristics are specified in **Supplementary Table 6**.

We studied 56,553 AF patients commenced on a DOAC (**Methods**). To predict time to all-cause mortality, we used baseline covariates, including choice of DOAC (from apixaban, rivaroxaban or dabigatran), age at onset of AF and age at DOAC initiation, sex, country of birth and ancestry, time from AF diagnosis to DOAC initiation, HAS-BLED^25^ and CHA2DS2-VASc^26^ scores, weight and BMI prior to DOAC initiation, and serum bilirubin, AST, ALT, ALP, platelet count, LDL, eGFR, creatinine, HbA1c, and haemoglobin prior to DOAC initiation. We excluded data from censored patients.

Time to all-cause mortality was available for 1,747 patients. Hyperparameter tuning (**Supplementary Fig. 1, Methods**) selected a PCHazard deep learning model^13^ with parameters detailed in **Supplementary Table 1**. As in use-case 2, we did not have a separate validation cohort available and so we performed nested cross-validation (**Methods**). Prediction accuracy was highest for shorter survival times which reflects the bias of the training data (**Fig. 5b**). Overall concordance was 0.59, with a median absolute difference between predicted and observed survival of 0.74 years with comparable performances for all DOACs. (**Fig. 5c, Supplementary Table 5**). We demonstrate that individualised prediction favours a different DOAC for specific patients (**Methods**, **Fig 5d, Supplementary Table 6**) who would not be differentiated by traditional guidelines.^27^

### Webpage implementation of predicTTE

predicTTE is a self-contained software package designed to facilitate the design and training of time-to-event prediction models. There is a case to be made for expanding access to trained models beyond those with the means to develop and train their own model. In a recent survey of ALS patients, 68.9% of patients expressed a preference to be able to access personalised data regarding their own survival prediction.^7^ This could be achieved through an online portal via which a trained model can be interrogated; we have produced this for the use-cases described in our manuscript (www.predictte.org**, Supplementary Video 1).** Another advantage of this online portal is that users, including clinicians and patients, could contribute their own data for model training (**Fig. 1**, **Supplementary Video 1**) and access our app to design and train their own models. We propose that our online portal offers some of the advantages of data collection via social media tools such as ease of use and accessibility, but without the challenges of trying to anonymise data that is freely available online;^28^ data submitted within the portal can be anonymised but is also securely stored with gold-standard SSL encryption.

## Discussion

Time-to-event prediction is relevant to the vast majority of human disease but there is a particular need for better predictions for Neurological diseases where tissue-based biomarkers are often missing. Improvements in prediction will likely provide accurate prognoses, guide personalised medicine and facilitate timely clinical interventions. However, rapid development in the statistical tools for time-to-event prediction has had limited impact because of the requirement for advanced statistical knowledge and computational expertise. Here we address this limitation through predicTTE, a new app and accompanying online portal (demonstrated in **Supplementary Video 1**). predicTTE provides cutting-edge computational tools to non-expert users who hold the appropriate clinical data. Moreover, our secure online portal enables sharing of data and distribution of individualised prediction to patients and clinicians, who can then contribute their own data via the online portal. We envision a positive feedback loop leading to exponential improvements in training data size and prediction model performance.

We present three use-cases for predicTTE where we have empirically addressed open questions in the field. We provide an implementation of a random forest model – MissForrest – for imputation of missing data. In testing this method, we considered whether the outcome variable should be used in imputation. We show that inclusion of the outcome variable can degrade model performance in a validation dataset. This is in opposition to previous findings^29^ but notably the contradictory study predates the development of the set of spline and deep learning time-to-event prediction tools we have applied. In particular, deep learning models may be more vulnerable to overfitting when missing values within the training data carry an artificial signature of the outcome variable.

In the prediction of ALS survival we also compared a single-step prediction model to an ensemble framework, whereby a final prediction is reached following a second stage of focused learning in a subset of the most informative patients within the training dataset; this is analogous to a transfer learning approach.^30^ We achieved a statistically significant improvement in model performance via this ensemble method. We hypothesise that this strategy will be most useful in future works focused on phenotypes such as ALS survival, where the distribution of events is significantly non-normal and skewed towards one extreme (**Fig. 2c**); most ALS patients die within five years but 10% can live longer than 10 years.^6^ Indeed we did not choose to apply the ensemble model configuration for the third use-case where survival of AF patients shows less skew (**Fig. 5b**).

In our three use-cases we have focused on the accuracy of an absolute time-to-event prediction rather than concordance or a range of probable values. In the context of ALS, this aligns with expressed patient preference.^7^ predicTTE can be used to develop prediction models where hyperparameter tuning and model performance is judged by concordance (**Supplementary Video 1**), and within the app and online portal, graphical outputs provide a probability distribution in addition to an absolute prediction value. In use-case 2, concordance and the absolute difference between predicted and observed survival were similarly altered between models. However, in use-case 1, concordance failed to capture the improvement in performance as a result of our ensemble model. A practical advantage of an exact prediction value is that results are optimally portable to other applications, as we demonstrate for discovery of genetic drivers of ALS survival.

A common approach which we have not applied is to use multiple imputation rounds where predictions are performed after each round of imputation so that uncertainty in the resulting prediction can be quantified via Rubin’s rules.^31^ This was not possible for our pipeline because Rubin’s rules assume a normal distribution of imputed estimates which is not true for the MissForrest model.

We have shown how predicTTE can be used in translational applications such as survival prediction, for the evaluation of covariates such as biomarkers, and even personalised treatment strategies. Our predicTTE framework is based on the idea that the key to seeing these applications become reality across many diseases, is the combination of optimal models with good quality large datasets.

## Disclosures

AAK is a consultant for NESTA. AAC reports consultancies or advisory boards for Amylyx, Apellis, Biogen, Brainstorm, Cytokinetics, GenieUs, GSK, Lilly, Mitsubishi Tanabe Pharma, Novartis, OrionPharma, Quralis, and Wave Pharmaceuticals.

## Acknowledgments

This work was supported by the National Institutes of Health (CEGS 5P50HG00773504, 1P50HL083800, 1R01HL101388, 1R01-HL122939, S10OD025212, P30DK116074, and UM1HG009442 to MPS), the Wellcome Trust (216596/Z/19/Z to JCK), NIHR (NF-SI-0617-10077 to PJS, 301648-RP to CJM, and NIHR202421 to AAC), the Motor Neurone Disease Association (MNDA) (899-792 to JCK, 975-799 to AAK and 943-793 to HM), the Swedish Research Council (2020-03485 and 2018-05973 to EE), the ALS Association (AAK), and the Darby Rimmer Foundation (AAK). We also acknowledge support from the NIHR Sheffield Biomedical Research Centre (NIHR203321) and the NIHR Sheffield Clinical Research Facility. We are very grateful to the ALS patients and control subjects who generously donated biosamples. We acknowledge transcriptomic data provided by the AnswerALS Consortium. Figures were created using BioRender.com.

## Methods

### Study Cohorts

#### Use-case 1

*Use of PrediTTE to predict survival in ALS*

#### Project MinE

The sporadic unrelated ALS patients included in this study as part of the Project MinE cohort were recruited at specialised neuromuscular centres in the UK, Belgium, Germany, Ireland, Italy, Spain, Turkey, the United States and the Netherlands.^14^ Patients were diagnosed with possible, probable or definite ALS according to the 1994 El-Escorial criteria.^32^ All controls were free of neuromuscular diseases and matched for age, sex and geographical location. The complete cohort consisted of 6,288 samples; 322 samples did not have a recorded observed or censored survival time and so were removed. Comparison of cohorts revealed atypically long survival in cohorts from Turkey and Portugal, as described elsewhere;^33^ these patients were removed to leave 5,336 patients for analysis.

#### UK validation cohort

661 patients who attended the Sheffield MND Care Centre were randomly selected. Patients received their diagnosis between 1999 and 2015. Baseline data were recorded from actual clinical reports and therefore this cohort is representative of real world data. Survival data were obtained or censored on 1^st^ August 2022. Survival was observed (not censored) in 595 (90%) of the patients; 179 (27%) of the patients had no missing data.

#### Use-case 2

*Survival prediction to evaluate candidate biomarkers in ALS*

We applied predicTTE to a previously published dataset consisting of 248 ALS patients with details of clinical symptoms at the time of biomarker sampling.^19^ Patients were diagnosed with possible, probable or definite ALS according to the 1994 El-Escorial criteria.^32^

#### Use-case 3

*Individualised prediction of all-cause mortality to guide preventative drug treatment*

*Clalit cohort:* Data from 56,553 patients with AF commenced on a DOAC were obtained from the centralised database of Clalit Health Services, Israel’s largest integrated healthcare provider and insurer. Established by the Clalit Research Institute, it serves more than half of Israel’s population of 4 million, with >95% of patients retained for five-years or more. A more detailed description of this dataset is available elsewhere.^24^

### Hyperparameter tuning and model choice

Model choice and hyperparameter tuning were guided by comparative testing of model configurations encompassing the full range of parameters within the pycox ^3^ implementation of MTLR,^34^ PC-Hazard,^13^ PMF,^13^ and DeepHit^35^ deep learning models. Tuned model parameters included layer structure, number of knots and learning rate (**Supplementary Table 1**). Similarly we tested forty flexible parametric models including the Royston-Parmar spline model, generalised gamma and generalised F-distributions as implemented within the flexsurv R package;^36^ the optimum spline model was implemented for comparison of prediction performance with the optimum deep learning model.

For each use-case, model selection utilised the total training set including imputed data and censored survival times. We divided the model choice process into a series of discrete steps to reduce the number of iterations and the potential for overfitting (**Supplementary Fig. 1**). Hyperparameter tuning was performed via nested cross-validation. We separated the data into 80% for training and 20%, which we designate the external validation dataset, for the final assessment of model performance. We then performed 10-fold cross validation within the 80% of data designated for training; here the data was *further* divided, on 10 separate occasions, into 80% for training and 20% for validation. We selected a random starting seed for each of the 10 rounds of cross-validation. Afterwards the 10 survival probability functions were summarised using their median values in order to achieve one uniformed prediction for evaluation in the external validation dataset.

In total hyperparameter tuning and model choice involves testing ∼2000 models (**Supplementary Figure 1**). Model evaluation is performed by combining three outcome measurements: concordance, the median absolute difference between actual and observed time to event, and the normalised median absolute difference (in which the absolute difference is divided by the observed time to account for deviations in short and long survivors equally). Models are ranked by performance in each outcome measure, and the lowest rank across all three measures is used to direct model choice (**Supplementary Fig. 2)**. predicTTE provides capacity to select alternative outcome measures in future use-cases.

In some instances we have compared model performance against a model trained using randomised covariate values. To facilitate a fair comparison we repeated hyperparameter tuning with the randomised data.

### Construction of an ensemble model

We developed an ensemble approach based on the hypothesis that patients with similar predicted time-to-event are more informative regarding the actual time-to-event of a test patient and, similarly, patients with very different time-to-event may be relatively uninformative (**Fig. 2A**). The ensemble model is constructed as follows: in the first step we trained the optimum deep learning model (determined by hyperparameter tuning) and used it to generate a prediction for a patient with unknown time-to-event. In the second step, the trained model is further trained with an additional 200 epochs utilising only the subset of the training cohort containing individuals with measured time-to-event within a range defined by the initial prediction value. The final model is then used to output a final predicted time-to-event.

We used the ensemble model in use-case 1 which concerned ALS survival. The distribution of ALS survival events is significantly non-normal and skewed towards shorter survivors (**Fig. 2c**) i.e. the majority of patients have shorter survival times. Therefore we used a dynamic range (± 70% of the predicted value) to select the focused training cohort used in the second stage of the ensemble model. The advantage of this approach is that the selected range of included survival times is larger for longer survival predictions than shorter survival predictions, such that the number of patients used in the focused training is approximately equivalent irrespective of the initial prediction. The +/-70% range was selected based on a level of accuracy that provided an improved prediction in four-fifths of the cohort.

### Training and testing optimal models

After model choice and hyperparameter tuning, the optimal model (determined by hyperparameter tuning) was trained using 10-fold cross validation. In each fold ∼80% of training samples were selected at random for model training and 20% of training samples are used for testing. After 10 folds, the final prediction is output as the median of the 10 different predictions. The C-index (Concordance) was calculated based on^37^ and in case of ties in predictions and event times adjusted according to^3^.

In use-case 1 each fold used the entire training dataset for model training because a separate independent validation dataset was available. In the other use-cases differences in training dataset size and the lack of an independent external validation cohort necessitated a nested cross-validation approach. In use-case 2, 5 samples were sequentially left-out from training and used for external validation of the trained model performance; in use-case 3, 10 samples were sequentially left-out from training and used for external validation of the trained model performance.

### Training and testing the MissForest model for imputation of missing data

A key aspect of our platform is the capability to impute missing data using a model called “MissForest”, which has shown superior performance in real-world testing^5^ and an ability to simultaneously handle continuous and categorical data.^4^ MissForest imputes data iteratively, starting with the variable with the least missing observations and progressing to the variable with the most missing observations. A random forest model is fit on the observed values. Each imputed value in this study relied on the mean result from 10 rounds of imputation because this represented the best compromise between computational time and independence of initial random seeds. In use-case 1, for testing predictions within the independent validation dataset, to avoid data leakage missing covariates for each patient were imputed using a new dataset including the training dataset and *only* the specific single patient from the validation dataset.

We validated the performance of the MissForest model using the 5,336 patients within the training dataset used for use-case 1, where the number of missing data ranged from site of onset (n=55) to ALSFRS-R slope (n=3,369 missing). In fifty different rounds we randomly selected and omitted 50 data points from each covariate in the training dataset. These data were then imputed using the MissForest model and the Pearson correlation between imputed and correct values was calculated.

### Identification of personalised DOAC treatment for AF patients

We sought to demonstrate, using our trained prediction model, that the optimal DOAC for an individual to maximise survival time may be impacted by a range of clinical factors. To investigate this we created a synthetic set of patients by dividing all continuous clinical covariates into quartiles and manufacturing all possible combinations of values. Prediction was performed on this synthetic dataset, determining the optimal DOAC for each patient. We demonstrate that the optimal DOAC is variable (**Fig. 5d**) between patients who would not be differentiated by traditional guidelines.^27^

### Software availability

The predicTTE online portal (https://www.predictte.org/) includes instructional material and links to download the app for different platforms.

## Supplementary Material Legends

**Supplementary Table 1: Hyperparameters for optimal prediction models in each use-case.**

**Supplementary Table 2: Prediction model performance for use-case 1 including randomised training data, and with/without imputation which included/did not include the outcome variable.** Model performance is measured by the difference, absolute difference, absolute difference normalised for patient survival and C-index. We also show the proportion of predictions within ’x’ years and % of actual patient survival. DL = optimum deep learning model. RP = Royston-Parmar which was the optimum spline model.

**Supplementary Table 3: Number (%) of missing data points for use-case 1**

**Supplementary Table 4: Prediction model performance for use-case 2.** Model performance is measured by the difference, absolute difference, absolute difference normalised for patient survival and C-index. We also show the proportion of predictions within ’x’ years and % of actual patient survival.

**Supplementary Table 5: Prediction model performance for use-case 3.** Model performance is measured by the difference, absolute difference, absolute difference normalised for patient survival and C-index. We also show the proportion of predictions within ’x’ years and % of actual patient survival.

**Supplementary Table 6: Example patients where choice of anticoagulation with either dabigatran, rivaroxaban, or apixaban impacts survival time.** Corresponding to **Fig. 5d**.

**Supplementary Video 1: Demonstration of the predicTTE online portal for model training, individualised prediction and data sharing**

**Supplementary Figure 1:**
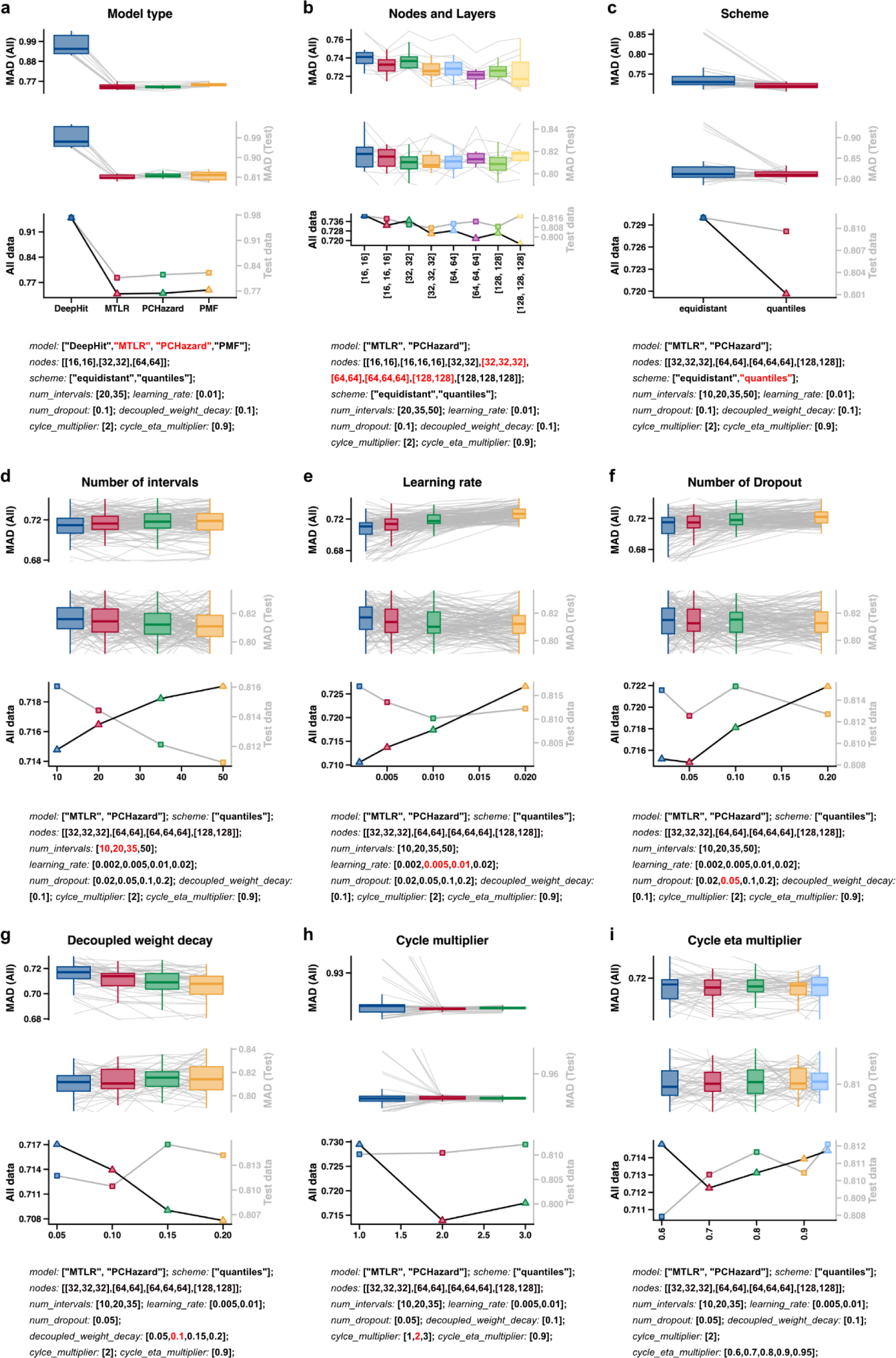
Hyperparameter tuning and model choice. The figure illustrates the step-by-step process of hyperparameter tuning. Plotted data denotes prediction performance derived from every tested combination of hyperparameters. Each panel demonstrates the effect of varying a single hyperparameter using the training data from use-case 1. The specific hyperparameter is noted in the title; boxplots are shown for the median absolute difference between actual and predicted time to event (MAD) in the entire training dataset (top subpanel), and in the external validation dataset (middle subpanel). For comparison, in the bottom subpanel the MAD value is shown for both the entire cohort (black line) and the external validation dataset (grey line). To avoid testing all possible combinations of hyperparameters, which could lead to overfitting, we select hyperparameters in a series of discrete steps. The first step includes hyperparameters which are more likely to have a large effect on prediction performance. In the first step only the *model type* was tested (**a**); in the second step the possible *nodes and layers* were reduced to a smaller subset and within the same models the *scheme* was selected (**b, c**). In the third step the *number of intervals*, the *learning rate* and the *number of dropout* were selected from the same model combinations (**d, e, f**). Finally the *decoupled weight decay* (**g**), *cycle multiplier* (**h**) and *cycle eta multiplier* (**i**) were chosen. Underneath each panel there is a list of all hyperparameters tested; red hyperparameters in a specific category are those taken forward to the next step of hyperparameter tuning whereas black hyperparameters in that same category are dismissed. The interpretation of hyperparameters is found in^3^. Other hyperparameters including the number of epochs and batch size had a minimal effect on prediction performance (data not shown).

**Supplementary Figure 2:**
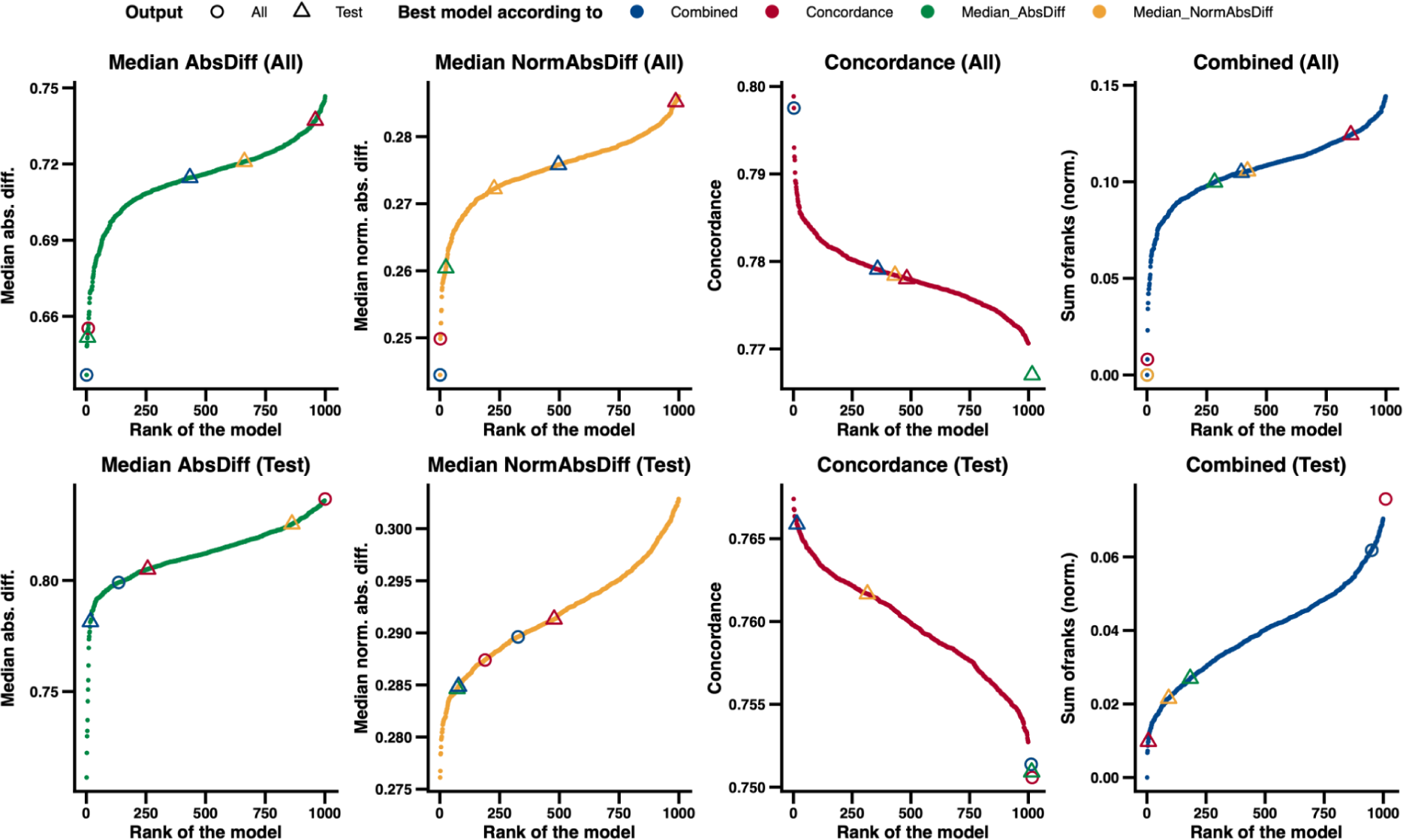
Ranking of models during hyperparameter tuning. This figure illustrates the concept behind choice of the optimal model via hyperparameter tuning. Model rank and performance relates to the best 1000 models tested during hyperparameter tuning for use-case 1, as described in **Supplementary Fig. 1**. The top row shows model rank and performance in the entire training set (60% training, 20% validation, 20% external validation) and the bottom row shows model rank and performance for the 20% external validation set. From left to right the outcome measurements: median absolute difference, normalised median absolute difference (median absolute difference divided by observed time), concordance and combined (normalised sum of the ranks of all three outcome measurements) are shown and coloured differently. For comparison, in each plot the rank of the best model according to a *different outcome measure* is shown in the corresponding colour, and with circles for the entire data and triangles for the external validation data. As demonstrated the best combined (summed rank for all outcome measures) model for the entire dataset (that was picked as the optimal model) performs well (is ranked low) in all other outcome measurements (blue circle) apart from the concordance in the external validation dataset, although the difference to the best model was low (0.015). On the other hand picking the model that performs best in the external validation set with respect to concordance (red triangle) leads to a relatively poor performance would be low in all other measurements.

**Supplementary Figure 3:**
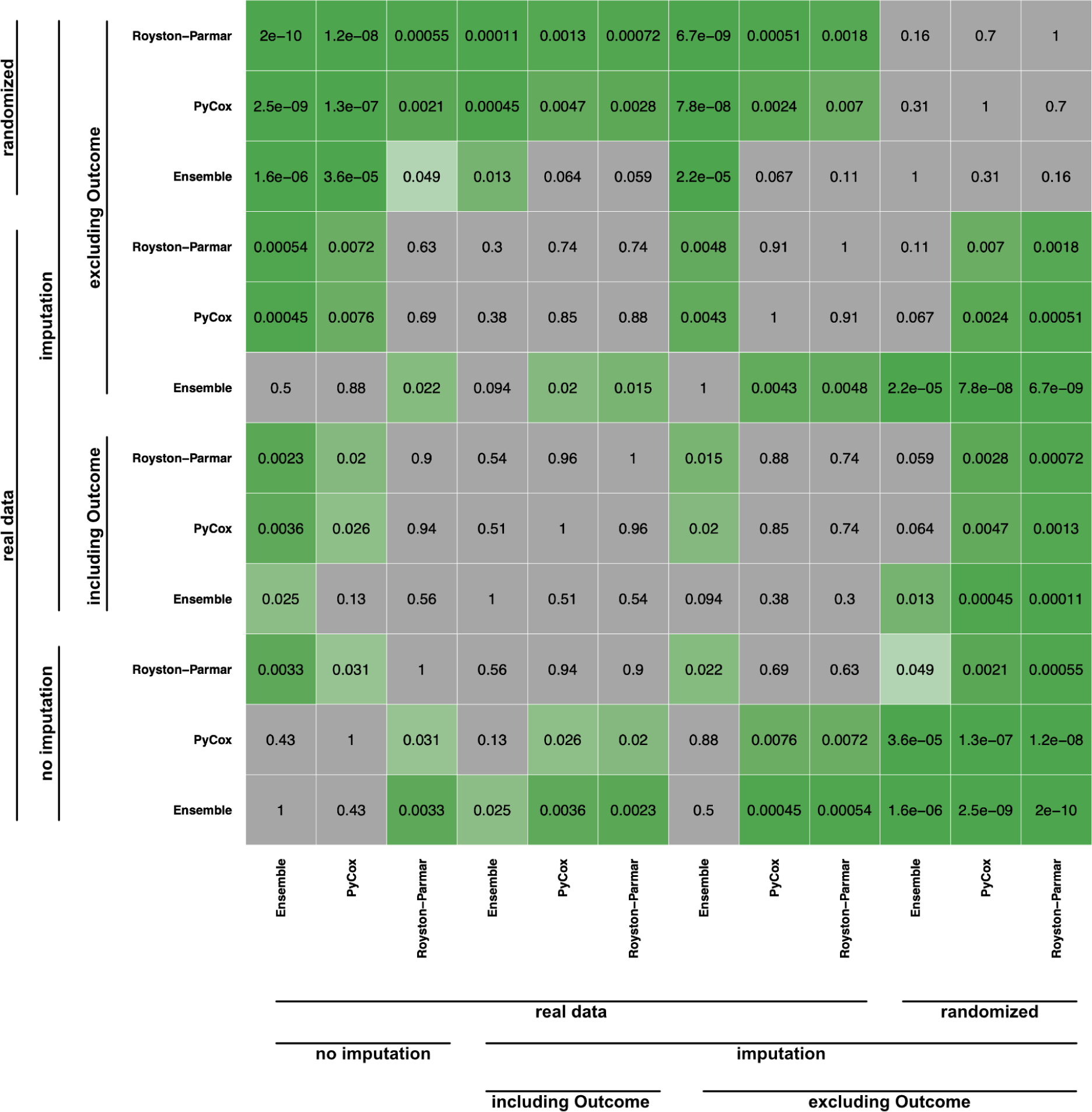
Prediction model performance for use-case 1 including shuffled training data, and with/without imputation which included/did not include the outcome variable. P-values are shown for a Wilcoxon rank-sum test comparing absolute difference between predicted and observed survival for each pair of models. Pycox = optimum deep learning model. Royston-Parmar = the optimum spline model. Ensemble indicates the ensemble model including a second stage of focused model training. The lowest median absolute difference between predicted and observed survival for a model which included imputation, was achieved with the ensemble model, with imputation which did not include the outcome variable (Fig. 3d); the difference between performance of this model and alternatives was statistically significant (p<0.05) with the exception of the ensemble model with imputation which *did* include the outcome variable (p=0.094).

## Notes

https://www.predictte.org

